# Maintenance of restriction-modification system diversity creates pan-immunity

**DOI:** 10.64898/2026.10.09.757998

**Authors:** Joseph Westley, Tatiana Dimitriu, Andrew C. Matthews, Iolanda Lopes Domingues, MultiDefence Consortium, Mark D. Szczelkun, Edze R. Westra, Stineke van Houte

## Abstract

Bacterial antiviral defense systems are highly diverse and unevenly distributed across populations, such that different strains carry distinct immune repertoires that together form a pan-immune system. Restriction-modification (RM) systems are the most abundant bacterial defense systems and exemplify this architecture, exhibiting remarkable diversity in structure, function, specificity, and efficacy. How and why such diversity is maintained, why weaker systems are not outcompeted, and whether diversity provides benefits beyond protection of individual cells remain unclear. Here, we experimentally tracked bacteria and phages across populations differing in RM diversity. We find no consistent evidence of a trade-off between defense strength and host fitness. Instead, we identify a trade-off between defense strength and horizontal transferability: stronger RM systems are less able to spread between strains, likely because factors that increase defense strength also increase toxicity risk upon transfer to unmethylated hosts. These weaker yet more mobile systems may therefore act as more accessible pan-immune components that bacterial populations can acquire in response to shifting parasite pressures. Furthermore, we show RM systems experience negative frequency-dependent selection, such that common specificities are more readily overcome by phages while rare ones remain protective, causing the benefits of potent systems to erode as they spread. Together, these dynamics maintain RM diversity and explain why weaker systems persist alongside stronger ones. Finally, we show that population-level RM diversity generates collective protection by constraining the emergence of RM-escaping phages and limiting the invasion of bacterial surface mutants that avoid phage adsorption, providing experimental support for the pan-immune model.

## Introduction

The arms race between prokaryotes and their viral parasites, bacteriophages (phages), shapes the structure of microbial communities (1) and drives the evolution of a diverse arsenal of defense mechanisms (2,3). It has been proposed that at the population level this defense system diversity can function as a collective resource, the pan-immune system, providing protection against phages while distributing the costs of defense system carriage (4). Of the >300 distinct defense system types described to date (5), the most abundant are restriction-modification (RM) systems, found in >80% of sequenced bacterial genomes (6). This abundance, combined with the broad protection they confer against phages (7–13) and other mobile elements (14–19), makes them a crucial component of the microbial pan-immune system. Although RM systems are mechanistically well characterized, the ecological and selective processes that maintain their diversity, and the consequences of this diversity for collective phage resistance, remain poorly understood.

Canonically, RM systems function by methylating self-DNA at a specific sequence and cleaving DNA if an unmethylated copy of the sequence is recognized (20). This allows discrimination between self– and non-self-DNA. Despite sharing this fundamental mechanism, RM systems vary in gene content, protein complex formation, methylation and cleavage mechanisms, and sequence specificity. The scale of this diversity is reflected in REBASE (a curated database of restriction enzymes and DNA methyltransferases), which as of 2022 listed >15,000 distinct active proteins, with over an order of magnitude more predicted from sequence data (21).

A key aspect of RM system functionality that varies considerably between systems is the strength of protection they confer against phages (7). *De novo* escape from RM commonly occurs when invading phage DNA is methylated by host methyltransferases before degradation by the cognate restriction enzymes. Variation in the frequency of this escape, together with differences in restriction enzyme efficiency, generates the substantial variability in defense strength. Among the systems tested to date, this has been shown to produce reductions in phage efficiency of plating (EOP; the relative ability of a phage to form plaques on a given host compared to a reference host) ranging from 1 to 8 orders of magnitude (7,8). This stark variation in defensive phenotype raises an evolutionary question: why are systems that confer weaker protection against phage not outcompeted by more effective alternatives? (7,8).

First, we hypothesize that defensive efficacy trades off against fitness cost. Traits enhancing RM defense, such as more efficient restriction or slower methylation of unmethylated DNA, also increase the risk of autoimmunity (22). Fluctuations in phage infection pressure could therefore favor weaker or stronger RM systems in different ecological contexts, allowing both to persist.

Second, we hypothesize that weaker RM systems are more readily acquired by new hosts. RM systems are highly mobile (23,24), frequently moving on mobile genetic elements (MGEs), but their acquisition is intrinsically risky because the recipient genome initially lacks the protective methylation signature. Mechanisms such as restriction alleviation can transiently reduce endonuclease activity until methylation is established (25), but this control is not universal. Where it is absent or imperfect, stronger RM systems should impose a greater risk of toxic autoimmunity during establishment, since the traits ensuring phage DNA is cleaved before it can be methylated will also act on a naive host genome. Weaker RM systems may trade protection for transmissibility, conferring less protection to hosts in which they are established but spreading more readily to new hosts. This transmissibility may allow bacterial populations to adapt more rapidly to encounters with novel phages.

Third, we hypothesize that negative frequency-dependent selection (NFDS) helps to maintain RM diversity. Although methylated escape phages can re-infect cells carrying the same RM specificity, this protection is lost upon replication in hosts with a different specificity, or none. Selection therefore favors escape phages adapted to the commonest RM variant, which both arise more frequently due to higher encounter rates and can subsequently infect a larger fraction of hosts (26). Dominant RM variants thus experience disproportionate phage pressure and decline in frequency, while rarer variants gain a relative advantage (27). Although theory suggests that NFDS maintains RM diversity (26), experimental evidence is lacking.

Our fourth hypothesis concerns the consequences of RM diversity rather than its maintenance. Diversity in resistance alleles is thought to limit the emergence and amplification of phages that overcome host resistance (28,29). Modeling predicts that RM specificities distributed across a population can render cells mutually uninfectable (30): escape phages produced by one cell carry a methylation pattern that does not protect them against other RM types, so infection is confined to strains carrying the same system. We therefore hypothesize that more RM-diverse bacterial populations will be collectively better protected against phages. This prediction, central to the pan-immune system framework (4), has not been tested empirically.

Using classical and full-spectral flow cytometry to track the relative densities of competing fluorescently labeled strains, we find no evidence that RM systems conferring weaker phage protection incur lower fitness costs, but they are more easily mobilized into new hosts. We show that phages can drive negative frequency-dependent selection among RM-carrying bacteria over the short term, and that higher population-level RM diversity limits the invasion of bacteria with alternative resistance strategies.

## Results

### Variation in RM protective benefits

To explore how population-level diversity is maintained when RM systems have varying effectiveness and cost, we first quantified the variation in protective benefits of three *Escherichia coli* derived RM systems (EcoRV, EcoRI, and EcoVIII).

*E. coli* strains carrying each RM system were exposed to a panel of diverse *E. coli* phages. These eight lytic phages (Table 1) represent seven distinct species and have a mean sequence similarity of 54.6% (Figure S1). EOP assays revealed substantial differences in the levels of protection conferred by each RM system even after controlling for counts of RM target sites in phage genomes. Specifically, EcoRV and EcoRI both reduced EOP similarly but significantly more than EcoVIII (*P* < 0.001, Figure 1A).

**Figure 1.**
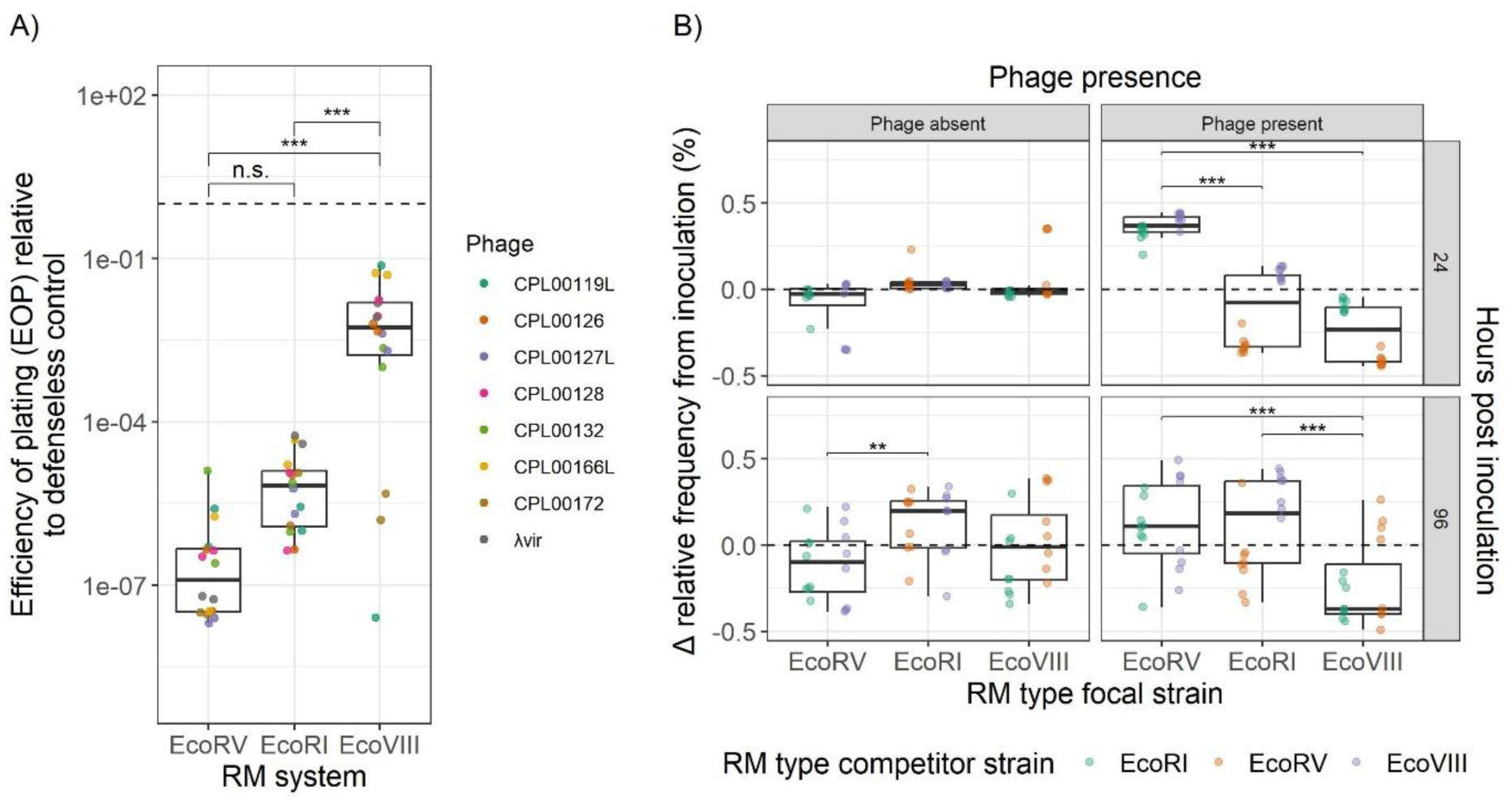
RM systems exhibit differences in phage resistance, which influence short term fitness in phage presence but do not consistently determine cost in phage absence. A) The efficiency of plating (EOP) of eight phages, on lawns of bacteria carrying one of three RM systems, relative to a control strain lacking the system. Points denote biological replicates (n = 2 per phage). Post-hoc pairwise comparisons are reported, indicating whether differences between RM systems remain statistically significant after controlling for the number of RM target sites in phage genomes (*** = *P* < 0.001, ** = *P* < 0.01, * = *P* < 0.05,. = *P* < 0.1). B) Change in relative frequency of an RM-carrying focal strain (RM type focal strain), competing with another RM-carrying strain (RM type competitor strain), from inoculation to 24 or 96 hours post inoculation, in the absence or presence of phage λvir. Points denote different biological replicates (n = 8 for each competing pair). Significance of post-hoc pairwise comparisons are included. For all boxplots central horizontal lines denote the median, boxes denote the interquartile range, and whiskers denote the most extreme values falling within 1.5 interquartile ranges of the box. Dashed horizontal lines denote neutral fitness. See Table S3 for specific post-hoc pairwise *P* values and effect size estimates for both figure panels A and B.

**Table 1.**
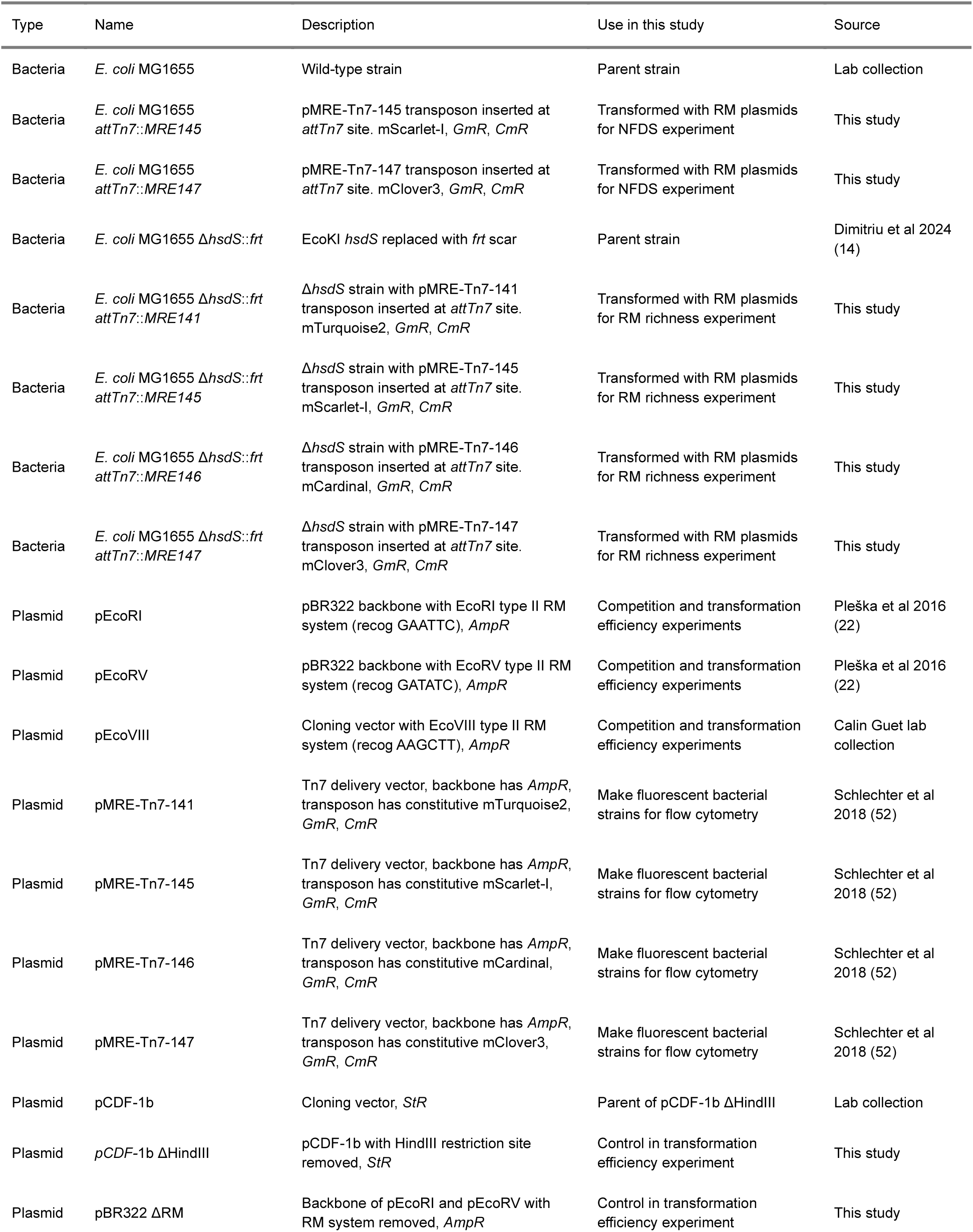
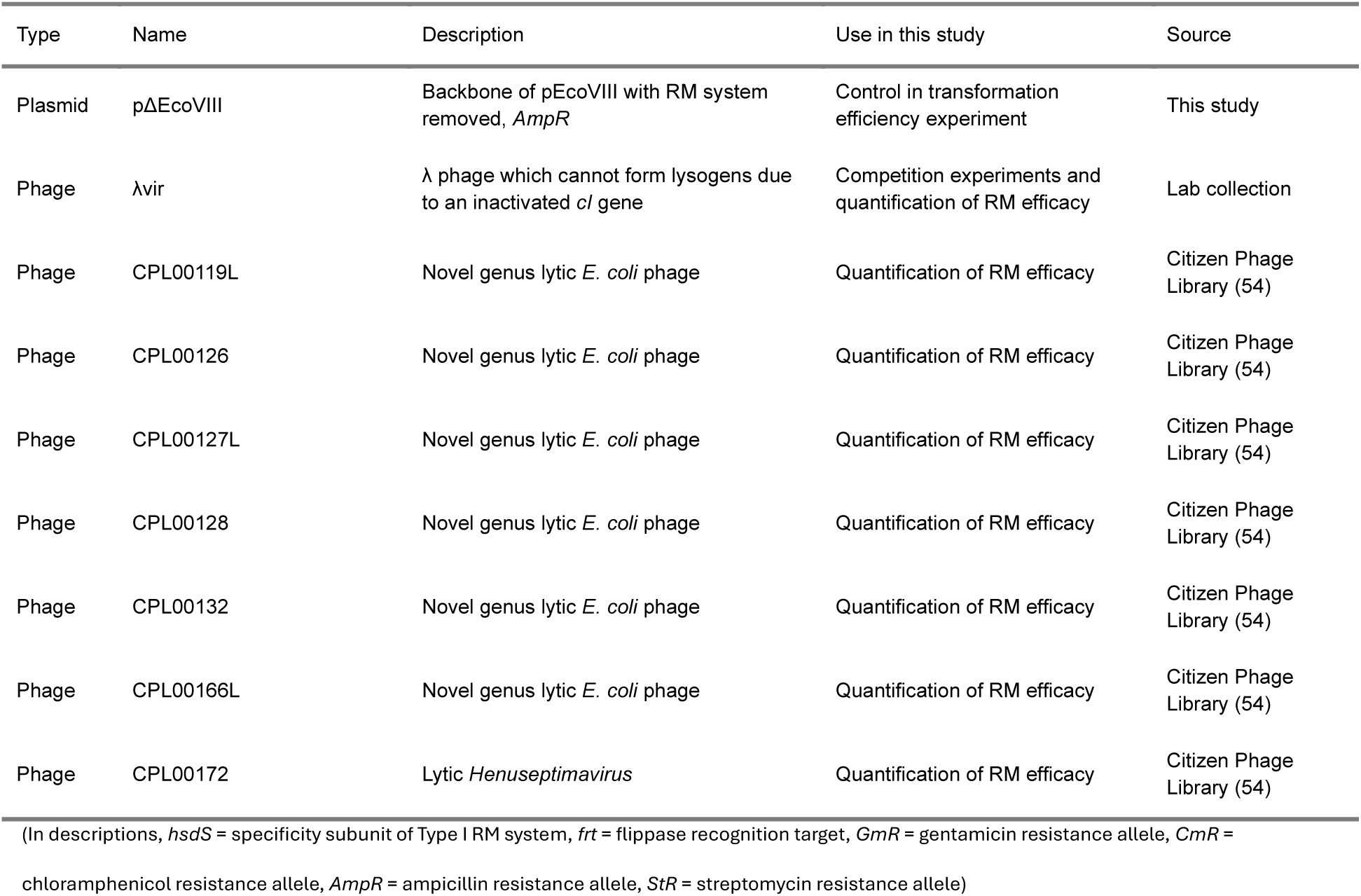
List of bacterial strains, plasmids, and phages used in this study.

To test how selection acts on these systems, we conducted all pairwise competitions in liquid media between fluorescently labeled versions of the RM-carrying strains. In the absence of phage, we found over the first 24 hours of the experiment no significant changes in relative frequency between strains carrying different RM systems (Figure 1B). When we considered the change in relative frequency over the full 96 hours of the experiment, we found that EcoRV (Δ relative frequency over 96 hours = –0.113) was significantly less competitive than EcoRI (Δ relative frequency over 96 hours = 0.114, Tukey-adjusted *P* = 0.002). However, although EcoRV was also less competitive than EcoVIII (Δ relative frequency over 96 hours = –0.008), this was not statistically significant, and neither was the difference between EcoRI and EcoVIII. These data show that in our experimental conditions, there is no evidence of a consistent trade-off between the level of phage resistance an RM system confers and its competitive ability in the absence of phage.

### The fitness advantage of stronger RM systems varies over time

Given variation in RM efficacy, we predicted that the more effective an RM system is against a phage, the more competitive it would be in the presence of that phage. To test this, we conducted pairwise competitions of strains carrying the RM systems in the presence of λvir, to which EcoRV confers the strongest protection (EOP < 5.9 × 10^-^⁸, the limit of detection), followed by EcoRI (EOP = 4.8 × 10^-5^), and finally EcoVIII (EOP = 1.2 × 10^-2^). These pairwise competitions revealed that EcoRV was significantly more competitive than EcoRI (Tukey-adjusted *P* < 0.001) and EcoVIII (Tukey-adjusted *P* < 0.001) at 24 hours post inoculation (Figure 1B). Consistent with our hypothesis that this is driven by variation in the protective benefits associated with the different RM systems, we observed the more strongly an RM system reduced λvir EOP, the more the strain carrying it increased in relative frequency (log10 EOP vs. change in relative frequency: R² = 0.682, F(1,46) = 101.9, P < 0.001). Notably, over longer timescales (96 hours), EcoRV was not consistently more competitive than EcoRI, although both were more competitive than EcoVIII (Figure 1B). This suggests that the relative fitness of the strains varies over time.

### RM systems that confer stronger defense are less easily mobilized to new hosts

The RM systems used in this study occur on mobile genetic elements (31–33), and we hypothesized that variation in their ability to transfer horizontally could contribute to the maintenance of defense diversity and the pan-immune repertoire. Specifically, we hypothesized that there might be trade-offs between defense efficacy and toxicity upon horizontal acquisition due to RM characteristics that are associated with weaker phage defense (faster methylation of unmethylated DNA and/or less efficient restriction) also reducing the risk of autoimmunity. To test this, we conducted an experiment where an *E. coli* host strain was simultaneously transformed with an RM-carrying plasmid (or empty plasmid backbone) and a control plasmid. The control plasmid lacked sites for EcoRV, EcoRI, and EcoVIII, to ensure RM systems could not restrict control plasmid uptake. From the relative transformation efficiencies of the RM containing plasmid and empty plasmid backbone, the reduction in transformation efficiency caused by RM presence was calculated.

We found that EcoVIII, the least effective defense system, did not reduce the transformation efficiency of the plasmid encoding it to the same extent (18.3%, 95% CI 5.5–31.1%) as the more effective defense systems, EcoRI (87.4%, 95% CI 74.6–100%; Tukey-adjusted *P* < 0.001) and EcoRV (71.8%, 95% CI 59.0–84.6%; Tukey-adjusted *P* < 0.001) (Figure 2). There was no statistically significant difference between the reduction in transformation efficiency of plasmids carrying EcoRI or EcoRV (Figure 2). These data demonstrate that RM systems that confer stronger defensive phenotypes can also be more toxic upon horizontal acquisition, suggestive of a defense efficacy-horizontal mobility trade-off.

**Figure 2.**
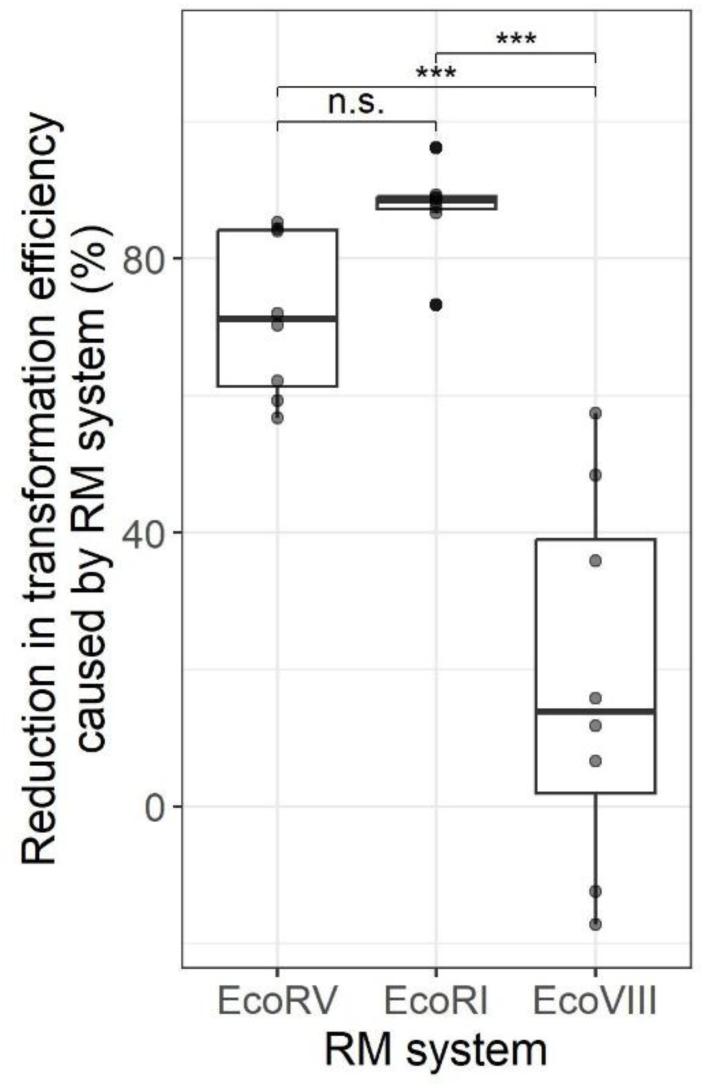
RM systems that confer higher phage resistance also cause higher reductions in the transformation efficiency of plasmids encoding them. Central horizontal lines denote the median, boxes denote the interquartile range, and whiskers denote the most extreme values falling within 1.5 interquartile ranges of the box. Points denote individual biological replicates (n = 8 per RM system). Significance of post-hoc pairwise comparisons are included (*** = *P* < 0.001, n.s. = *P* > 0.1). See Table S3 for specific post-hoc pairwise *P* values and effect size estimates.

### Phages drive negative frequency-dependent selection for RM systems

To investigate if phages can drive negative frequency dependence of bacteria carrying RM systems (26,34), we conducted pairwise competitions with varying starting frequencies of the RM-carrying strains (5:95, 50:50, 95:5). Each strain in a pair is plotted in turn as the *focal* strain against its *competitor* (Figure 3), so ratios refer to the focal strain’s starting frequency: rare, equal, or common. In the absence of phage, strains maintained their starting frequency as expected from their similar cost (above). However, in the presence of phage, the change in relative frequency was negatively associated with the starting frequency of the bacterial strains: rarer strains were more competitive than more common strains (Figure 3). Although the strength of this frequency dependence (the slope) was similar across strains and competitions, strains still differed in their baseline fitness. The strain carrying EcoRV had the highest baseline fitness, regardless of its competitor. In contrast, the EcoRI– and EcoVIII-carrying strains showed lower fitness when competing against EcoRV but had similar fitness when competing against each other. Despite these differences in baseline fitness, phages drove strong negative frequency-dependent selection for RM systems in every competition.

**Figure 3.**
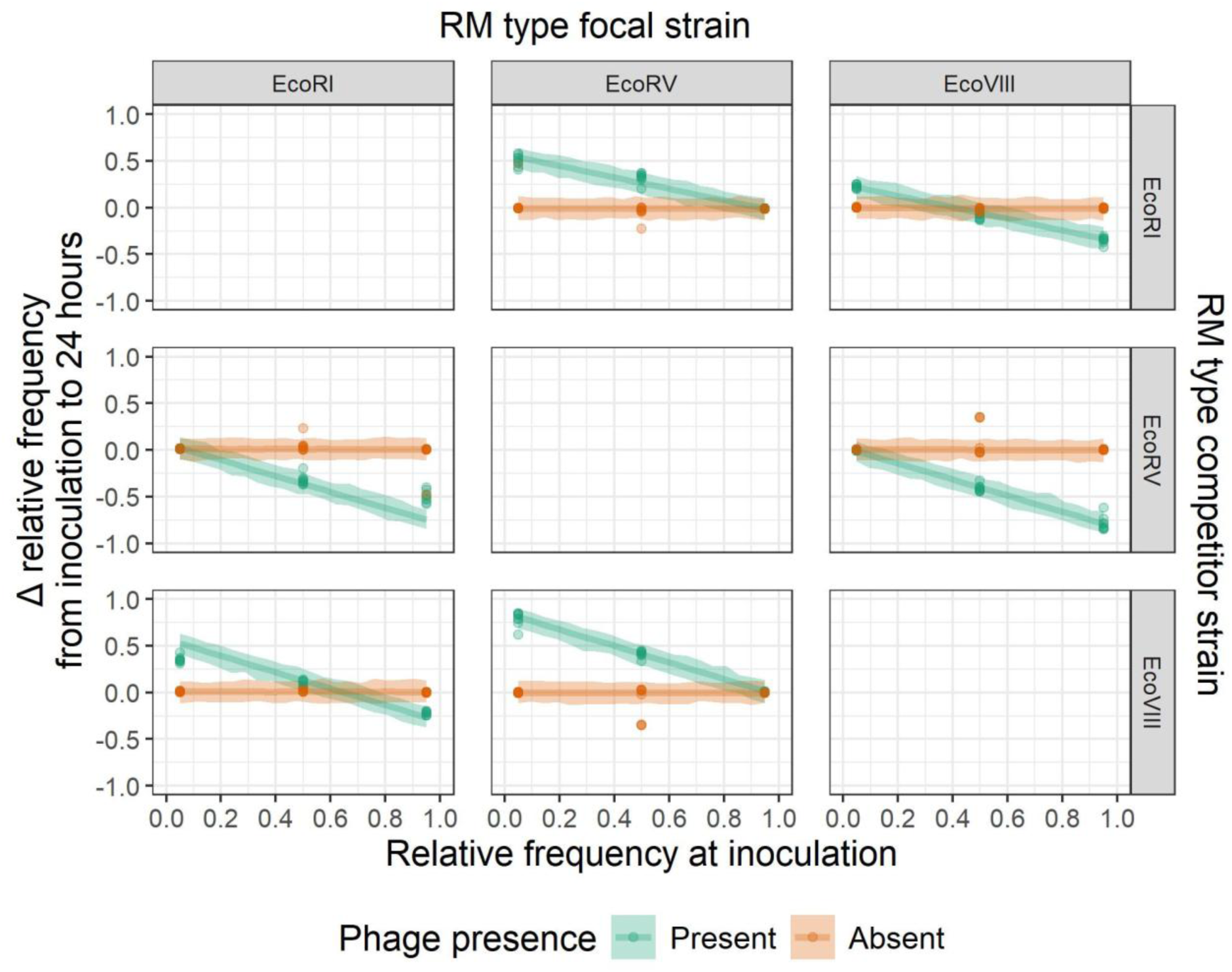
Phages drive negative frequency-dependent selection of RM systems. The relationship between strain frequency at inoculation and its change in frequency from inoculation to 24 hours, in the presence and absence of phage. Column labels denote the RM system of the focal strain, and row labels denote the RM system of the competitor strain. Individual points denote biological replicates (n = 8 per treatment group), lines denote predictions, and ribbons 95% credibility intervals.

Notably, negative frequency-dependent selection was transient; from 24 to 48 hours post inoculation, we observed weakly positive frequency-dependent selection in the presence of phage (Figure S2), and from 48–96 hours relative frequencies remained stable (Figure S3B). Both the weak positive frequency dependence and subsequent stability arose due to the invasion of phage resistant surface mutants (Figure 4); surface mutants most often emerged from the dominant strain at 24 hours (as the dominant strain experiences the highest mutation supply), then invaded and remained at high frequency (Figure S3B). These data show that frequency-dependent selection contributes to the maintenance of RM diversity until surface mutants invade.

**Figure 4.**
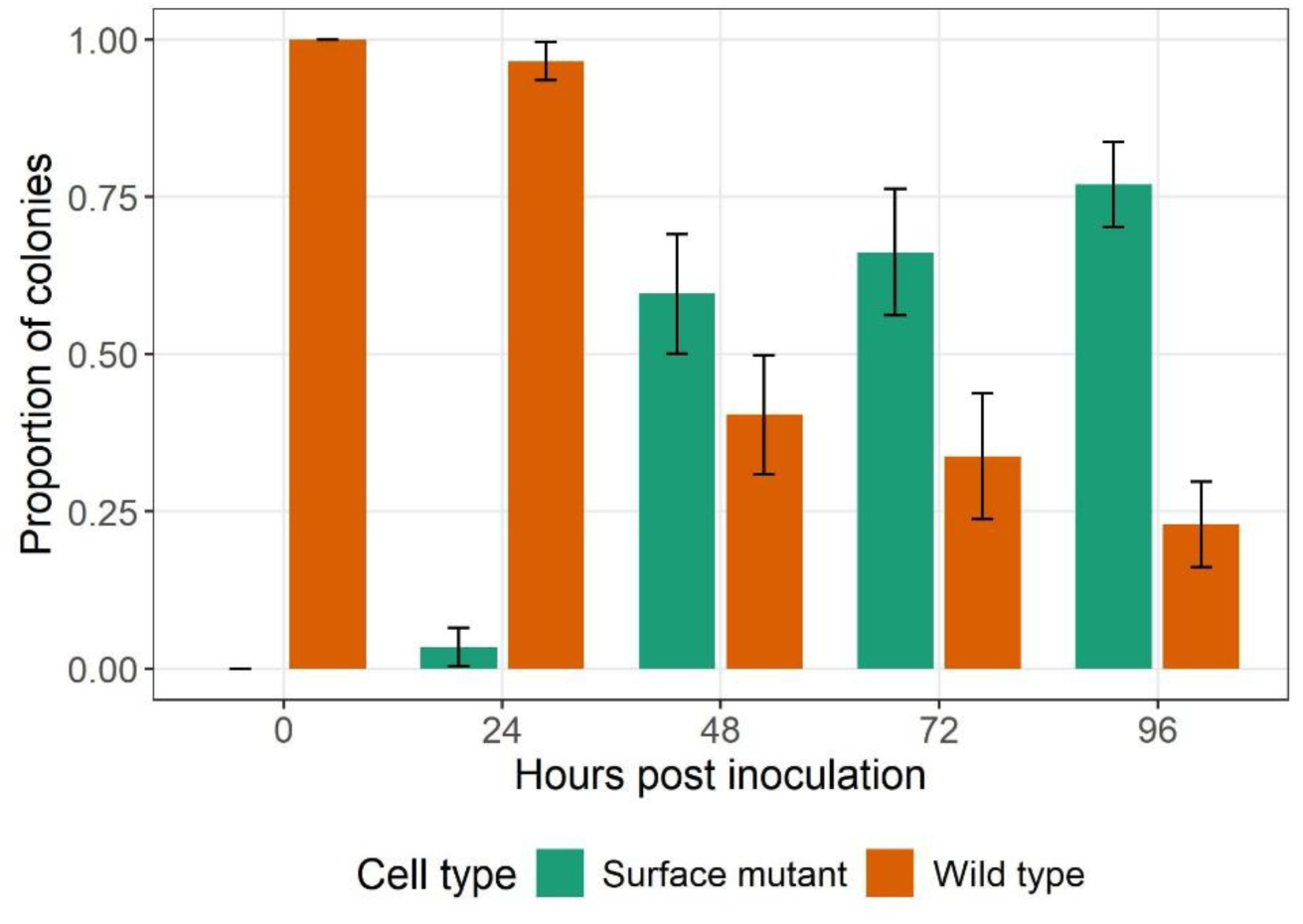
Surface mutants increase in relative frequency over time. The relative frequencies of surface mutant (maltose auxotrophic) cells that are resistant to λvir at each timepoint for all competition experiments combined. Error bars indicate 95% confidence intervals of the mean proportion of colonies.

### Higher population level RM diversity reduces force of infection and slows invasion by surface mutants

We next examined how RM diversity affects bacteria–phage coevolution. To test whether more diverse RM populations provide stronger protection against phage infection, we varied RM richness, defined as the number of distinct RM-carrying subpopulations present in a culture, from zero to three. Treatments included monocultures of each RM-carrying strain and an RM-negative control, all pairwise combinations of RM-carrying strains, and a mixture containing all three RM-carrying strains. We predicted that, if RM diversity strengthens population-level protection against λvir, surface receptor mutants should invade more slowly as RM richness increases, due to reduced selection for alternate resistance strategies. Consistent with this prediction, higher RM richness slowed the increase in the proportion of surface mutants over time (likelihood ratio test: χ²(9) = 68.77, *P* < 0.001; Figure 5A). Notably, the RM system that was most effective against λvir was not consistently dominant in pairwise or three-way competitions at 96 hours (Figure S4). Because all phage-treated populations were ultimately dominated by surface-resistant mutants, this likely reflects stochastic differences in the timing of resistance mutations rather than differences in RM efficacy.

**Figure 5.**
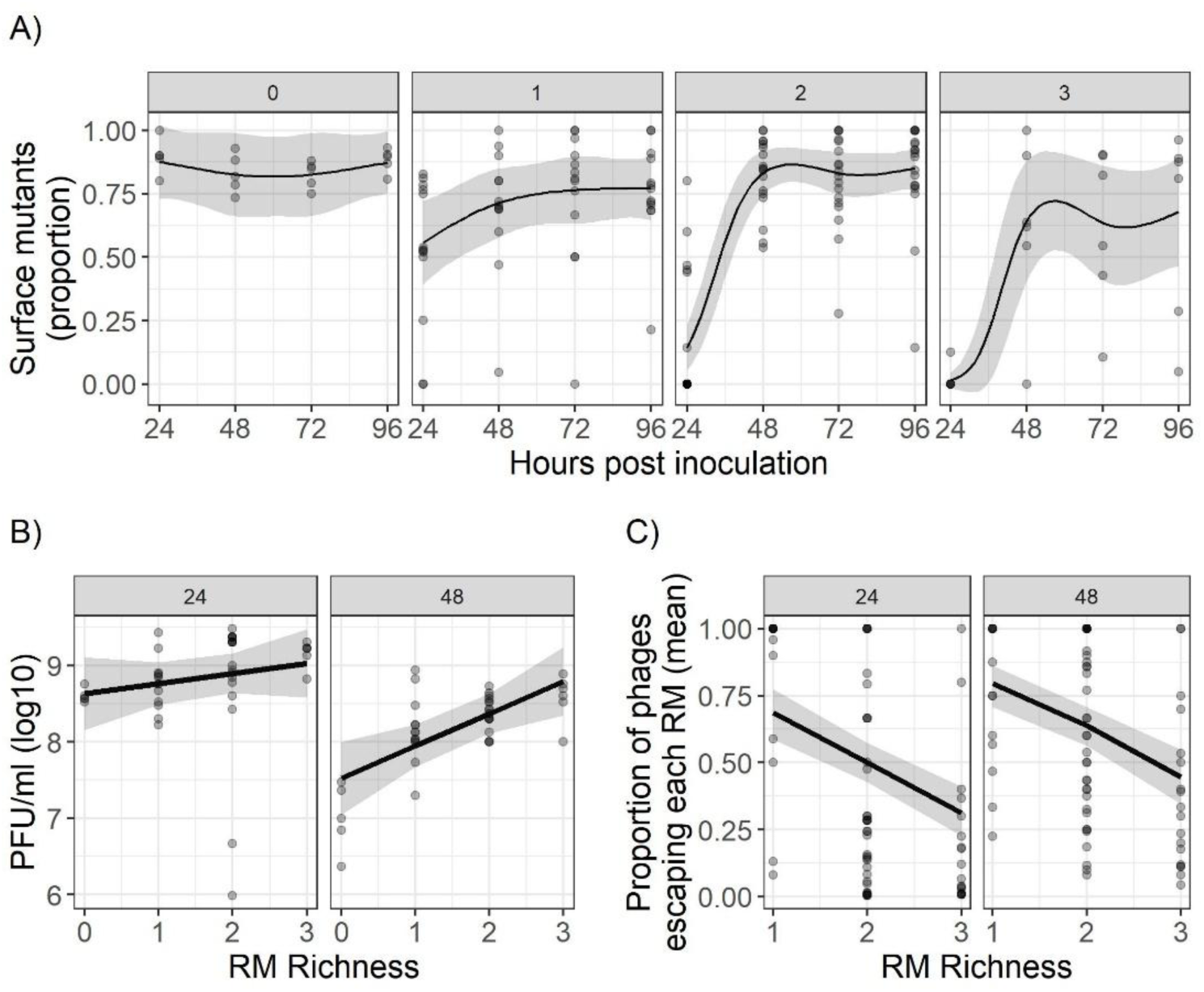
Higher population level RM system diversity reduces the force of infection and slows invasion by surface mutants. A) The proportion of total cells that are λvir resistant surface mutants across time. Panels indicate treatments with varying RM system richness. B) The relationship between RM richness and the total plaque forming units per ml (PFU/ml (log10)), for 24 and 48 hours post inoculation. C) The mean proportion of total phages escaping each RM system present in the respective treatment group for 24 and 48 hours post inoculation. For all panels, points indicate individual biological replicates (n = 6 for each possible combination of respective RM richness), solid black lines denote the relationship as predicted by generalized linear mixed models (GLMMs) and ribbons denote the prediction’s 95% confidence interval.

We next asked whether the slower invasion of surface receptor mutants in more diverse RM populations could be explained by changes in phage abundance and the infection pressure experienced by each RM-carrying subpopulation. We therefore quantified both total phage titer and, for each RM system, the proportion of phages able to escape restriction. Contrary to our initial expectation, total phage titer increased with RM richness, and this relationship strengthened between 24 and 48 hours post inoculation (likelihood ratio test: χ²(1) = 5.78, P = 0.016; Figure 5B). This likely reflects the rapid invasion of surface receptor mutants in low-richness treatments (Figure 5A), which reduced the density of susceptible hosts and consequently limited phage amplification. In contrast, the proportion of phages able to escape a given RM system declined significantly with RM richness (likelihood ratio test: χ²(1) = 22.33, *P* < 0.001; Figure 5C). Thus, although more diverse RM populations supported larger phage populations overall, each RM-carrying subpopulation was exposed to fewer phages capable of infecting it. These results support the pan-immunity prediction that defense diversity can protect individual bacterial lineages by reducing their effective force of infection.

Notably, phage extinction was observed by 24 hours in five out of six replicates in which EcoRV-carrying strains were exposed to λvir in monoculture; these replicates were excluded from subsequent analyses. Phage extinction was not observed in any other treatment. This reveals an important consequence of defense diversity: although diverse RM communities can reduce the effective force of infection experienced by individual subpopulations, they may also prevent phage extinction by providing more weakly defended hosts that sustain phage replication. Thus, pan-immunity does not necessarily mean that all strains benefit equally from defense diversity. Instead, collective protection may emerge alongside strain-level trade-offs, where diversity lowers RM-specific infection pressure but can also sustain phage populations that would otherwise be lost.

## Discussion

Over recent years, the repertoire of known microbial defense systems has expanded rapidly (35,36), yet RM systems remain the most abundant defenses employed by prokaryotes (6). Here, we elucidate the mechanisms that allow for RM diversity to be maintained, and how this shapes population-level resistance to phage infection.

We find no evidence that RM systems conferring stronger resistance are more costly in the absence of phage. The autoimmunity costs of RM may be conditional on stress, as repair of stress-induced DNA damage can produce unmodified sites that are susceptible to self-cleavage (37). We did not impose such stress in our high resource growth conditions. However, we find that the strength of resistance associated with RM systems does trade off with their ability to mobilize to a new host. This mobility-resistance trade-off may have broader implications for the evolution of pan-immunity, which relies on frequent HGT of defenses to generate diversity and enable adaptation under heterogeneous parasite pressure (4). Our data suggest that RM systems that offer weaker protection may be more mobile and therefore play a more pronounced role in the early adaptation of species in a microbial community to novel phages. It will be informative to explore how selection for RM systems with different strengths encoded by MGEs depends on the mobility of the MGE (38–40).

We show that RM diversity can be maintained through negative frequency-dependent selection, but that this was transient and followed by weakly positive frequency-dependent selection due to the invasion of surface mutants (and these mutants are more likely to arise in the most abundant genotype). Surface mutants resist λvir through mutation of the LamB receptor at minimal fitness cost under our experimental conditions. In natural systems, however, where resources are limited and interbacterial competition is intense, surface mutation is likely to carry a greater cost (41–43), potentially prolonging phage-driven NFDS and the maintenance of RM diversity.

The observed NFDS in our experimental populations represents the underlying dynamic that drives fluctuating selection, where alleles fluctuate in their adaptive value over time (27,44). NFDS of RM systems likely arises because phage escape is epigenetic (and non-heritable when propagating on a strain without the escaped RM), preventing phages from accumulating methylation patterns that facilitate escape of all RM specificities in a population. This mechanism implies that phage driven NFDS may occur for other defenses where escape depends on non-heritable acquisition of host-specific signatures (such as BREX (45), DISARM (46), and MADS (47)). NFDS can also arise where heritable escape (antidefense systems) trades off against host range. In natural *Vibrio* populations that show patterns consistent with fluctuating selection, phages alter their host range by exchanging antidefense genes (48). Although accumulation of all antidefenses would allow a broader host range, the authors hypothesize that strict limits on phage genome size preclude this (49). Phage-driven NFDS may therefore be a widespread and enduring feature of natural microbial communities where phage escape of host defenses relies on epigenetic modification by host machinery, or where other factors constrain bacteria and phages from continually evolving increased resistance and infectivity (49,50).

More broadly, NFDS of RM systems may help maintain strain diversity. Because phages carry the methylation signature of their previous host, the most common strain absorbs most of the infection pressure. Mathematical modeling has predicted that this can sustain the coexistence of many lineages carrying different RM systems (30,34), even when they vary in their competitive ability in the absence of phage (34). Selection acts on entire lineages rather than on defense loci alone, so phage–defense coevolution may shape community structure and sustain genetic and functional diversity well beyond defense systems themselves.

Beyond identifying mechanisms that may maintain RM diversity, our results show that increasing RM richness reduces the effective force of infection and slows the spread of alternative resistance strategies. This provides empirical support for the pan-immune hypothesis that population-level defense diversity can collectively protect bacterial communities from phage infection (4). In this respect, RM diversity parallels CRISPR-Cas spacer diversity, where genetically distinct resistant subpopulations collectively limit phage spread and constrain the emergence of broadly infective phage mutants (28,29). More broadly, these findings echo work in eukaryotic host–parasite systems showing that immune diversity can enhance population-level protection against parasites (reviewed in (51)).

However, this collective benefit was not shared uniformly across strains. In particular, the strongest RM system could drive rapid phage extinction in monoculture but lost this extinction benefit in more diverse communities where less well protected strains sustained phage replication, even when they were rare. Thus, RM diversity is consistent with the pan-immune hypothesis but also reveals a strain-level trade-off: RM diversity can protect the community by reducing RM-specific infection pressure, while simultaneously sustaining phage populations that disadvantage the most strongly defended strains. More broadly, our findings suggest that immune diversity can generate collective protection while redistributing parasite pressure across genetically distinct host lineages.

## Materials and methods

### Strains and growth conditions

For all bacterial strains, plasmids, and phages used in this study see Table 1. Unless otherwise stated, all liquid cultures were grown in lysogeny broth (LB) at 37 °C and shaken at 180 RPM, and all solid media cultures were grown statically overnight at 37 °C on LB agar plates (1.5% agar). Where antibiotics are used for plasmid maintenance or selective plating, concentrations are as follows: ampicillin, 100 µg/ml; chloramphenicol, 25 µg/ml; streptomycin, 50 µg/ml.

### Producing RM-carrying fluorescent *E. coli* MG1655 strains

In all experiments, we used either *E. coli* MG1655, or an *E. coli* MG1655 strain with the native EcoKI RM system inactivated (MG1655 *ΔhsdS::frt*; see Dimitriu et al 2024 (14) for cloning of this strain). Transposons carrying fluorophore-encoding genes were inserted into the MG1655 chromosome through Tn7 transposon delivery plasmids (52). Chemically competent *E. coli* MG1655 cells (53) were transformed with the pMRE-Tn7 plasmids (see Table 1 for plasmids used). Transformed cells were grown overnight at 30 °C in 6 ml of media supplemented with 0.1% arabinose to induce expression of the transposase. The following day 60 µl of culture was transferred to 6 ml of fresh media without arabinose and was grown overnight at 42 °C (pMRE-Tn7 plasmids are conditional suicide plasmids that do not replicate above 37 °C). Fresh media was inoculated and grown again at 42 °C as one passage was found to be insufficient for plasmid loss. Cultures were plated and grown overnight on plates supplemented with chloramphenicol to select for cells carrying the transposon. The following day colonies were picked and streaked onto both plates supplemented with ampicillin and plates supplemented with chloramphenicol. Successful transposon acquisition and plasmid loss were determined when growth was observed on the chloramphenicol plate (indicating transposon acquisition) but not the ampicillin plate (indicating plasmid backbone loss). These fluorescent strains of MG1655 were then made chemically competent (53) and transformed with plasmids carrying one of three Type II RM systems (EcoRI, EcoRV, or EcoVIII, see Table 1).

### Assaying RM efficacy against a diverse phage panel

#### Phage panel

We tested the efficacy of RM systems against a panel of eight lytic *E. coli* phages (Table 1).

These phages were kindly provided by the Citizen Phage Library (54), and were chosen based on several criteria, including being listed as a lytic *Escherichia* phage, access to a complete genome sequence, the presence of at least two recognition motifs for all three RM systems used in this study in their genomes, and the ability to replicate in our *E. coli* MG1655 strain. Seven phages from the CPL met all these conditions, as did the phage λvir already present in our laboratory collection, providing us with a panel of eight lytic phages.

To quantify the genetic diversity of this panel of phages, we conducted multiple sequence alignment using MAFFT (v7.490) (55) with the FFT-NS-2 algorithm and default parameters (gap open penalty = 1.53, offset value = 0.123, scoring matrix 200PAM / k=2).

#### Phage amplification

Phages were amplified by spotting serial dilutions of existing phage stocks (provided by the CPL) onto lawns of *E. coli* MG1655 and incubating overnight. The following day, for each phage, 6 ml of media was inoculated with 60 µl of overnight grown culture of *E. coli* MG1655, grown for 30 minutes, and then inoculated from a single plaque picked from the streaked phage plate, and grown overnight. The following day lysed cultures were filtered through 0.22 µm 25 mm PTFE Syringe Filters to produce phage lysate. Resulting phage concentrations ranged from 1.3 × 10^9^ to 2.7 × 10^10^ PFU/ml.

#### RM system EOP assay

Serial dilutions of phage stocks were prepared in M9 media, and 3 µl of each dilution was plaque assayed (56) on lawns of *E. coli* MG1655, as well as lawns of *E. coli* MG1655 transformed with each of the RM-carrying plasmids (pEcoRI, pEcoRV, and pEcoVIII).

To quantify the defensive efficacy of each RM system, we calculated the efficiency of plating (EOP) for each phage on the lawn carrying the system, compared with the MG1655 control lawn. When no plaques were observed at any dilution on an RM lawn, EOP values were determined based on the limit of detection of the assay. Note that although *E. coli* MG1655 has the chromosomally encoded EcoKI RM system, as all phages were amplified on *E. coli* MG1655, they were premethylated by the host EcoKI methyltransferase and were thus resistant.

Therefore, here we were testing only the EOP reduction caused by the plasmid encoded RM systems. This was confirmed by plating *E. coli* MG1655 amplified phages on both wild type *E. coli* MG1655, and the EcoKI knockout strain MG1655 Δ*hsdS*::*frt*, and observing no difference in estimated PFU/ml.

### Competition and coevolution experiments overview

Competition and coevolution experiments were conducted in deep 96-well plates covered with Breathe-Easy breathable membranes to ensure sterility but allow gas exchange, in 1 ml of liquid media. All treatments were initially inoculated with a total volume of 10 µl of culture grown overnight in glass vials. All plasmids carrying RM systems and ampicillin resistance genes were maintained through addition of ampicillin to media. All cultures were transferred into fresh media at 24, 48, and 72 hours (10 µl of culture into 990 µl of fresh media). Samples were taken every 24 hours post inoculation, with final sampling at 96 hours. From these samples, bacterial strain titers were determined through flow cytometry (see Materials and methods: Flow cytometry data collection and analysis), phage titers were assayed through plaque assay (see Materials and methods: Plaque assay), and the proportion of cells that had lost the ability to metabolize maltose through a suspected mutation in the *lamB* gene that confers resistance to λvir phage (57) was assayed through use of indicator plates supplemented with maltose and Triphenyl Tetrazolium Chloride (TTC) (see Materials and methods: Surface mutant detection assay).

### Experimental design: Negative frequency dependence of RM systems

We conducted pairwise competitions between *E. coli* MG1655 strains carrying one of three RM systems encoding plasmids with the relative starting frequencies of 5:95, 50:50, and 95:5. Competitions in the presence and absence of phage λvir were conducted, with an inoculating multiplicity of infection (MOI) of 0.1. An MOI of 0.1 was chosen as pilot experimental investigation showed this allowed maintenance of phages and bacterial cells at 24 hours. To distinguish between the two competing bacterial strains with flow cytometry, we used the strains *E. coli* MG1655 *attTn7*::*MRE145* (producing the mScarlet-I fluorophore) and *E. coli* MG1655 *attTn7*::*MRE147* (producing the mClover3 fluorophore). Reciprocal competitions were conducted where the strains were switched, to control for any differential fitness effects caused by production of different fluorophores. Eight biological replicates of every competition were conducted. Cell titers were assayed using the Attune NxT flow cytometer (see Materials and methods: Flow cytometry data collection and analysis). Growth conditions and sampling protocols were as outlined in Materials and methods: Competition and coevolution experiments overview.

### Experimental design: Population-level RM system diversity and surface mutant invasion

We varied population-level RM system diversity across treatments from zero to three RM systems. In multi-strain treatments starting frequencies were kept equal, and total volume of inoculum was kept equal between all treatments. Competitions in the presence and absence of phage λvir were conducted, utilizing an inoculating MOI of 0.1. Six biological replicates of each treatment were conducted. Bacterial strains used in this experiment were: *E. coli* MG1655 Δ*hsdS*::*frt attTn7*::*MRE141* (producing mTurquoise2 fluorophore), *E. coli* MG1655 Δ*hsdS*::*frt attTn7*::*MRE145* (producing mScarlet-I fluorophore), *E. coli* MG1655 Δ*hsdS*::*frt attTn7*::*MRE146* (producing mCardinal fluorophore), and *E. coli* MG1655 Δ*hsdS*::*frt attTn7*::*MRE147* (producing mClover3 fluorophore). Cell titers were assayed using the Cytek Aurora spectral flow cytometer (see Materials and methods: Flow cytometry data collection and analysis). Growth conditions and sampling protocols were as outlined in Materials and methods: Competition and coevolution experiments overview.

*E. coli* MG1655 Δ*hsdS*::*frt* strains were used in this experiment (compared with *E. coli* MG1655 in the negative frequency dependence of RM systems experiment) as originally the design incorporated EcoKI as an additional experimental RM system; however, this was not pursued. In all experiments phages were propagated on the respective host strain so the presence or absence of EcoKI did not affect phage resistance.

### Experimental design: RM system horizontal mobility

To compare the transformation efficiencies of each RM-carrying plasmid, competent MG1655 Δ*hsdS*::*frt* cells were transformed with a mixture of test and control plasmids, plated onto selective plates, and the relative transformation efficiency of test and control plasmids was calculated. Five test plasmids were used: the three RM plasmids (pEcoRI, pEcoRV, pEcoVIII), and two empty vectors from which the RM systems had been removed (pBR322 ΔRM (this backbone is shared for EcoRI and EcoRV) and pΔEcoVIII). The control plasmid used to standardize the transformation efficiencies of the test plasmids against was pCDF-1b ΔHindIII (pCDF-1b with the EcoVIII restriction site removed to prevent the EcoVIII RM system interfering with transformation of the control plasmid, note pCDF-1b has no EcoRI or EcoRV restriction sites). All plasmids were propagated on MG1655 Δ*hsdS::frt*. Chemically competent MG1655 Δ*hsdS::frt* cells were produced using transformation and storage solution (TSS) buffer, as outlined in Chung & Miller 1993 (53), and stored in 30 µl aliquots at –70 °C. Competent cells were then thawed on ice for 20 minutes. For each transformation a mixture of 1 fmol of both test and control plasmid, in 1 µl of elution buffer, was added to the competent cells and incubated for 20 minutes on ice. Transformations were submerged in 42 °C water for 50 seconds, then cooled on ice for 2 minutes. 120 µl of LB was added and transformations were incubated at 37 °C and shaken at 180 RPM for 45 minutes. Transformations were plated onto both streptomycin and ampicillin supplemented plates, that were incubated overnight and then colonies were counted. We calculated the transformation efficiency ratio for each transformation as the number of cells transformed with the test (RM-positive or RM-negative) plasmid divided by the number transformed with the control plasmid. To quantify the percentage change in transformation efficiency caused by RM system presence, we normalized the ratio of the RM-positive plasmid transformations to the mean ratio for the corresponding RM-negative transformations, and expressed the result as a percentage change:

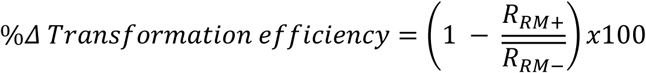

where *R*_RM+_ is the transformation efficiency ratio for an RM-positive plasmid, and *R_RM_*_−_ is the mean transformation efficiency ratio for the corresponding RM-negative plasmid.

### Flow cytometry data collection and analysis

#### Negative frequency dependence of RM systems

Cultures were sampled at 24, 48, 72, and 96 hours, paraformaldehyde was added to a final concentration of 4% to fix the cells, samples were agitated for 20 minutes on a microplate shaker and then diluted 1000-fold with phosphate buffered saline (PBS). Diluted samples, single-color controls, non-fluorescent wild type controls, and PBS only were analyzed on a four-laser Attune NxT flow cytometer equipped with an autosampler (Thermo Fisher Scientific, USA), at a flow rate of 25 µl/min, and a sampled volume of 50 µl. For each sample, a minimum of 5000 events gated as bacterial cells were recorded (the mean actual events gated as bacterial cells across samples was ∼30,000).

#### Population-level RM system diversity and surface mutant invasion

Cultures were sampled at 24, 48, 72, and 96 hours, and diluted 1000-fold with PBS (samples were not fixed with paraformaldehyde, as laboratory protocol deemed it unnecessary for the instrument used). Diluted samples, single-color controls, non-fluorescent wild type controls, and PBS only were analyzed on a five-laser Cytek Aurora full-spectrum flow cytometer (Cytek, USA), at a flow rate of 10 µl/min, and a sampled volume of 10 µl. For each sample, a minimum of 2000 events gated as bacterial cells were recorded (the mean actual events gated as bacterial cells across samples was ∼20,000).

#### Analysis in FCS Express

Cytometry data were analyzed with FCS Express 7 Research Edition. For both experiments, bacteria were gated to separate cells from debris with the forward scatter (FSC) and side scatter (SSC) signals, through comparison of PBS only, and cell containing samples. Fluorescence gates were drawn based on inspection of all single-color controls.

For Attune NxT collected data, all events that were gated as cells were plotted onto a BL1 and YL2 plot, where mScarlet-I and mClover3 cells were gated (Figure S5). No unmixing or compensation was required. For Cytek Aurora collected data, unmixing was conducted using the inbuilt Unmixing Wizard of FCS Express and the single-color and non-fluorescent controls. Autofluorescence compensation was not required. Detectors for gating were selected based on which detector overlapped with the peak of the respective fluorophore’s emission spectrum and will hereafter be referred to by the fluorophore’s name. All events that were initially gated as cells were plotted onto an mClover3-mScarlet-I plot, where mClover3 and mScarlet-I cells were gated. All events from this plot that were not gated were plotted onto an mScarlet-I-mCardinal plot, where mCardinal cells were gated. Finally, all ungated events from this plot were plotted on an mOrange-mTurquoise2 plot, where mTurquoise2 cells were gated (see Figure S6 for an overview of this sequential gating).

### Surface mutant detection assay

The phage λ infects *E. coli* through adsorption to the outer membrane protein, LamB, that facilitates diffusion of maltose into the cell. *E. coli* commonly evolves resistance to λ phage through mutations in the *lamB* gene, which frequently leads to loss of the cell’s ability to metabolize maltose (57). To identify LamB surface mutants, we prepared serial dilutions of culture samples in M9 media and spotted them onto LB agar plates supplemented with 0.005% Triphenyl Tetrazolium Chloride (TTC) and 1% maltose. Here, surface mutants that cannot metabolize maltose grow as red colonies and cells with functional *lamB* genes grow as white colonies (58). This protocol was verified through plating of the *lamB* knockout from the Keio collection (JW3996) (59) on these indicator plates and observing red colony growth. However, the assay does not capture all resistant cells: a random subset of white, maltose-metabolizing colonies proved resistant to λvir, including to phages that had escaped the RM system of the colony in question (likely through acquisition of its protective methylation signature). Across replicates, ∼80% of colonies with full resistance to λ phage grew as red, while ∼20% were morphologically indistinguishable from sensitive colonies. This is consistent with previous work showing that *E. coli* can acquire mutations in surface proteins that confer resistance to λ phage without compromising the ability to metabolize maltose (57). Due to this, the assay underestimates the true proportion of mutants with surface modification-based phage resistance in the population (i.e. when ∼80% of colonies are red on our TTC and maltose supplemented indicator plates (averaged across replicates) the true proportion of cells that have acquired a mutation conferring resistance to λvir is probably closer to 100%).

### Plaque assay

In both competition experiments, phages were purified from samples using chloroform extractions at each timepoint. Serial dilutions of phage extractions were prepared in M9 media, and plaque assayed (56) on lawns of *E. coli* MG1655 Δ*hsdS::frt*, as well as lawns of each RM-carrying strain. From this we could calculate the total phage titer in each replicate at each timepoint, as well as the titer of phages that had escaped each RM system.

### Statistical analysis

All statistical analysis was conducted in R v. 4.3.1 (60). The *tidyverse* packages (61) were used for data handling. To create figures the R packages *ggplot2* (61), *ggpubr* (62), *cowplot* (63), *RColorBrewer* (64), *gridExtra* (65), and *patchwork* (66) were used. To create tables *flextable* (67) and *ztable* (68) were used. The R package *brms* was used to conduct all Bayesian statistical analyses (69). All frequentist statistical analyses utilized Generalized Linear Models (GLMs), which were built using either the base R function *glm*, or the package *glmmTMB* (70). The package *DHARMa* (71) was used to validate GLMs.

#### Bayesian statistical methods

For all Bayesian models we used 4 chains, 20000 iterations, a warmup of 4000, a thinning interval of 10, a max tree depth of 15, and target acceptance probability for the No-U-Turn Sampler (NUTS) was set at 0.95. Chain convergence was confirmed through visual inspection of trace plots, ensuring R-hat was below 1.1, and effective sample sizes were >1000. We conducted prior predictive checks to ensure that the priors were uninformative yet did not yield implausible predictions; posterior predictive checks to confirm that the models could reproduce data consistent with the observed data; prior sensitivity analyses to verify that inferences were robust to reasonable changes in prior specifications; K-fold validation (K = 5), using the function *brms::kfold* to check predictive performance (ensuring the standard error of the expected log predictive density (elpd_kfold) was relatively low in comparison to the elpd_kfold).

For the models investigating how relative frequency of a strain at timepoint X impacts change in relative frequency over the following 24 hours, we used a Bayesian model with family specified as *student*. The response variable was the change in frequency between timepoint X and timepoint X+24 hours, the fixed effects modeled included the frequency at timepoint X, focal RM system identity, phage presence, the RM system identity of the competitor, all two-way interactions between these fixed effects, and three-way interactions between RM identity of the competitor, frequency at timepoint X, and focal RM identity, and RM identity of the competitor, frequency at timepoint X, and phage presence. Additionally, we included the fluorophore type as a fixed effect to control for differential fluorophore fitness costs.

#### Frequentist statistical methods

All frequentist statistical analyses were conducted using generalized linear models (GLMs) or generalized linear mixed models (GLMMs). For each response variable, an initial full model was constructed including all relevant explanatory variables and interactions, guided by the experimental design and biological hypotheses (Table S1). Significance testing was performed via model simplification. Terms were retained in the minimal adequate model if their removal significantly reduced model fit (*P* < 0.05). Model assumptions were checked using the *DHARMa* package, which uses simulation-based residual diagnostics. We tested for overdispersion, model fit (using a Kolmogorov-Smirnov test), influential outliers, and heteroscedasticity of residuals. See Table S1 for specific model families and fixed effect structures, and Table S2 for statistical significance of all fixed effects and interaction terms.

## Resource availability

### Lead contact

Requests for further information and resources should be directed to and will be fulfilled by the lead contact, Joseph Westley.

### Materials availability

Genetically engineered fluorescent *E. coli* strains are available upon request. No unique plasmids were generated for this manuscript.

### Data and code availability

Upon acceptance of the manuscript for publication all data and code used will be deposited at Zenodo and will be publicly available via a DOI, as of the date of publication. Prior to acceptance, all data and code will be made available to reviewers via inclusion as a zipped directory in the submission.

## Acknowledgments

This work was funded by a grant from the European Research Council to M.D.S. (ERC-2017-ADG-788405) and a BBSRC sLoLa grant to E.R.W. (BB/X003051/1). S.v.H. acknowledges funding from the UK Government’s Horizon Europe funding guarantee (EP/X026507/1). T.D. is supported by a Royal Society University Research Fellowship (URF\R1\231740). J.W. was supported by the BBSRC-funded South West Biosciences Doctoral Training Partnership (training grant reference DTP3: BB/T008741/1).

We thank Ben Temperton (University of Exeter) for providing several of the phages used in this study, and feedback on the manuscript, Claudia Igler for providing feedback on the manuscript, Exeter Centre for Cytomics for flow cytometry technical support, and Calin Guet for providing the RM system plasmids used in this study.

## Author contributions

**Westley** – conceptualization, investigation, methodology, formal analysis, data curation, visualization, writing – original draft, writing – review and editing

**Dimitriu** – conceptualization, investigation, methodology, supervision, writing – review and editing

**Matthews** – investigation, methodology, writing – review and editing

**Lopes Domingues** – investigation, methodology, writing – review and editing

**Szczelkun** – supervision, funding acquisition, writing – review and editing

**Westra** – conceptualization, supervision, funding acquisition, resource provision, writing – review and editing

**van Houte** – conceptualization, supervision, funding acquisition, resource provision, writing – review and editing

## Declaration of generative AI and AI-assisted technologies in the writing process

During the preparation of this work, the authors used Claude selectively to improve the clarity of the manuscript. After using Claude, the authors reviewed and edited the content and take full responsibility for the content of the published article.

## Supplemental information titles and legends

**Document S1.** Supplemental tables and figures (Tables S1-S3, Figures S1-S6).

