## Supplementary material for "Maintenance of restriction-modification system diversity creates pan-immunity": Document S1. Supplemental tables and figures (Tables S1-S3, Figures S1-S6).

**Table S1.** Overview of frequentist GLMs and GLMMs including full fixed effect structure prior to model simplification, random effect structures, and any potential model fit issues as identified by DHARMa (model validation notes).

| Figure | Response variable | Full fixed effect structure | Random effect structure | Model validation notes | Model family |
|---|---|---|---|---|---|
| Figure 1A | Log10 EOP relative to strain lacking respective RM | RM type * count of RM sites in phage genome | (1\|phage identity) | Slight quantile deviations (heteroscedastic residuals) | Gaussian |
| Figure 1B | Δ relative frequency of strain carrying an RM system | RM type * phage presence * timepoint | | | Gaussian |
| Figure 1B | Δ relative frequency of strain carrying an RM system | log10 λvir EOP relative to strain lacking respective RM | | | Gaussian |
| Figure 2 | Transformation efficiency reduction (%) | RM type | | | Gaussian |
| Figure 5A | Maltose auxotroph surface mutant cells (%) | RM richness * ns(timepoint, df = 3) | (1\|treatment/ replicate) | | Betabinomial |
| Figure 5B | log10 plaque forming units | RM richness * timepoint | (1\|treatment/ replicate) | | Gaussian |
| Figure 5C | Mean prop of phages escaping each RM in population | RM richness * timepoint | (1\|treatment/ replicate) | K.S. test significant | Beta |

**Table S2.** Significance of fixed effects as tested through model simplification for all GLMs and GLMMs. Note, where interaction terms are significant constituent fixed effects are retained in minimal model.

| Figure | Response variable | Fixed effect tested | *P* value |
|---|---|---|---|
| Figure 1A | Log10 EOP relative to strain lacking respective RM | RM_type:count of RM sites in phage genome | 0.868 |
| Figure 1A | Log10 EOP relative to strain lacking respective RM | RM_type | < 0.001 |
| Figure 1A | Log10 EOP relative to strain lacking respective RM | Count of RM sites in phage genome | < 0.001 |
| Figure 1B | Δ relative frequency of strain carrying an RM system | log10 λvir EOP relative to strain lacking respective RM | < 0.001 |
| Figure 1B | Δ relative frequency of strain carrying an RM system | RM type:phage presence:timepoint | 0.015 |
| Figure 2 | Transformation efficiency reduction (%) | RM type | < 0.001 |
| Figure 5A | Maltose auxotroph surface mutant cells (%) | RM richness:ns(timepoint, df = 3) | < 0.001 |
| Figure 5B | log10 plaque forming units | RM richness:timepoint | 0.016 |
| Figure 5C | Mean prop of phages escaping each RM in population | RM richness:timepoint | 0.148 |
| Figure 5C | Mean prop of phages escaping each RM in population | RM richness | < 0.001 |
| Figure 5C | Mean prop of phages escaping each RM in population | timepoint | 0.010 |

**Table S3.** Statistical significance and effect sizes for all post-hoc pairwise comparisons relating to Figure panel 1A, 1B, and Figure 2. Comparison indicates which RM pair are being compared, Effect estimate indicates the predicted effect size on the scale of the respective response variable (positive values indicate that the former RM system in Comparison has a higher value than the latter), Lower and Upper 95% CI indicate the lower and upper 95% confidence intervals of the predicted effect, *P* value indicates the Tukey adjusted *P* value, and Statistical significance is defined as: *** = $P < 0.001$, ** = $P < 0.01$, * = $P < 0.05$, . = $P < 0.1$.

| Figure panel | Comparison | Effect estimate | Lower 95% CI | Upper 95% CI | *P* value | Statistical significance |
|---|---|---|---|---|---|---|
| 1A | EcoRV - EcoRI | -0.127 | -1.68 | 1.42 | 0.980 | n.s. |
| 1A | EcoRV - EcoVIII | -2.18 | -3.64 | -0.713 | 0.001 | ** |
| 1A | EcoRI - EcoVIII | -2.05 | -3.17 | -0.936 | < 0.001 | *** |
| 1B, phage absent, 24 hours | EcoRI - EcoRV | 0.121 | -0.038 | 0.280 | 0.174 | n.s. |
| 1B, phage absent, 24 hours | EcoRI - EcoVIII | -0.014 | -0.172 | 0.145 | 0.977 | n.s. |
| 1B, phage absent, 24 hours | EcoRV - EcoVIII | -0.135 | -0.294 | 0.024 | 0.115 | n.s. |
| 1B, phage present, 24 hours | EcoRI - EcoRV | -0.481 | -0.641 | -0.322 | < 0.001 | *** |
| 1B, phage present, 24 hours | EcoRI - EcoVIII | 0.137 | -0.022 | 0.296 | 0.108 | n.s. |
| 1B, phage present, 24 hours | EcoRV - EcoVIII | 0.618 | 0.459 | 0.777 | < 0.001 | *** |
| 1B, phage absent, 96 hours | EcoRI - EcoRV | 0.228 | 0.069 | 0.387 | 0.002 | ** |
| 1B, phage absent, 96 hours | EcoRI - EcoVIII | 0.116 | -0.044 | 0.275 | 0.202 | n.s. |
| 1B, phage absent, 96 hours | EcoRV - EcoVIII | -0.112 | -0.271 | 0.047 | 0.220 | n.s. |
| 1B, phage present, 96 hours | EcoRI - EcoRV | 0.003 | -0.156 | 0.161 | 0.999 | n.s. |
| 1B, phage present, 96 hours | EcoRI - EcoVIII | 0.352 | 0.193 | 0.511 | < 0.001 | *** |
| 1B, phage present, 96 hours | EcoRV - EcoVIII | 0.349 | 0.190 | 0.508 | < 0.001 | *** |

| Figure panel | Comparison | Effect estimate | Lower 95% CI | Upper 95% CI | *P* value | Statistical significance |
|---|---|---|---|---|---|---|
| 2 | EcoRI - EcoRV | 15.6 | -6.38 | 37.6 | 0.197 | n.s. |
| 2 | EcoRI - EcoVIII | 69.1 | 47.1 | 91.1 | < 0.001 | *** |
| 2 | EcoRV - EcoVIII | 53.5 | 31.5 | 75.5 | < 0.001 | *** |

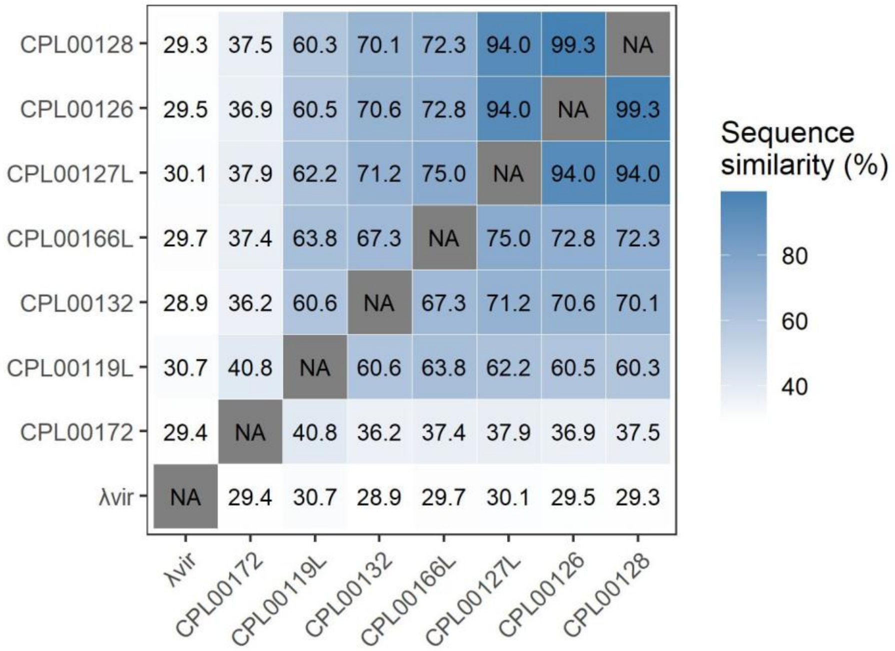


**Figure S1.** Pairwise sequence similarities (%) of phage genomes as calculated using MAFFT (v7.490) (55).

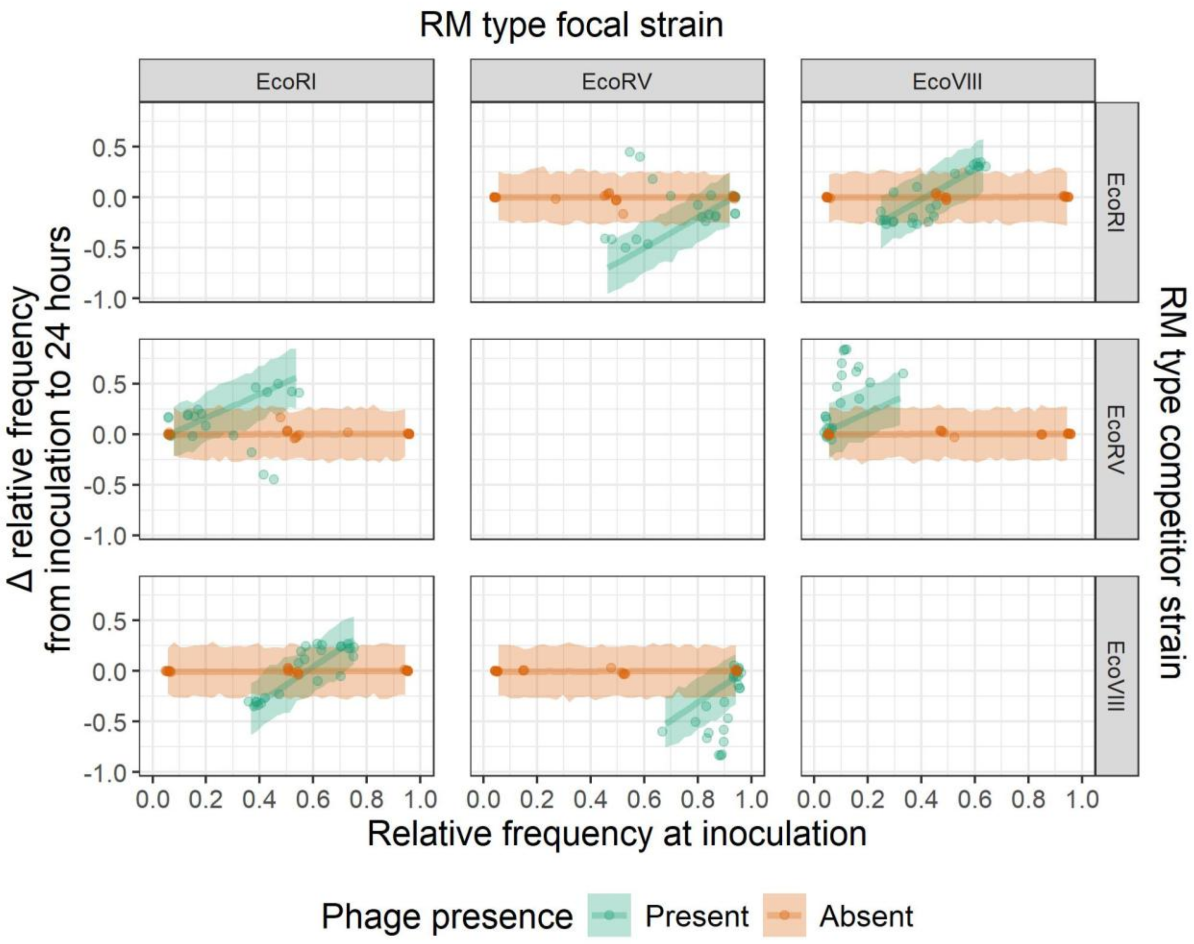


**Figure S2**. **Phages no longer drive negative frequency-dependent selection from 24-48 hours post inoculation.** The relationship between strain frequency at 24 hours and its change in frequency from 24 to 48 hours, in the presence and absence of phage. Column labels denote the RM system of the focal strain, and row labels denote the RM system of the competitor strain. Individual points denote biological replicates (n = 8 per treatment group), lines denote predictions, and ribbons 95% credibility intervals.

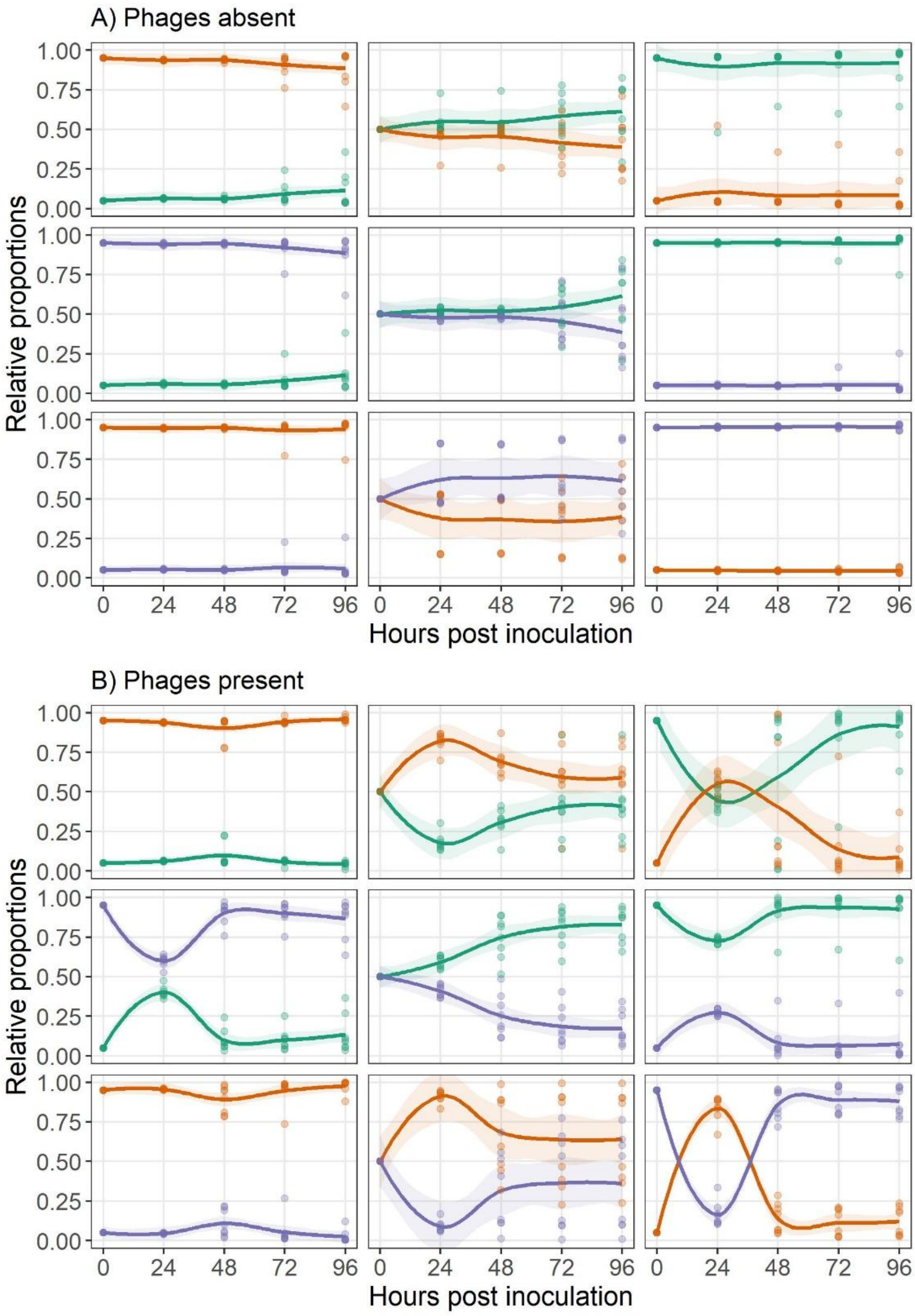


**Figure S3.** The change in relative frequencies of competing RM carrying strains over the course of the experiment in the absence (A) and presence (B) of phages.

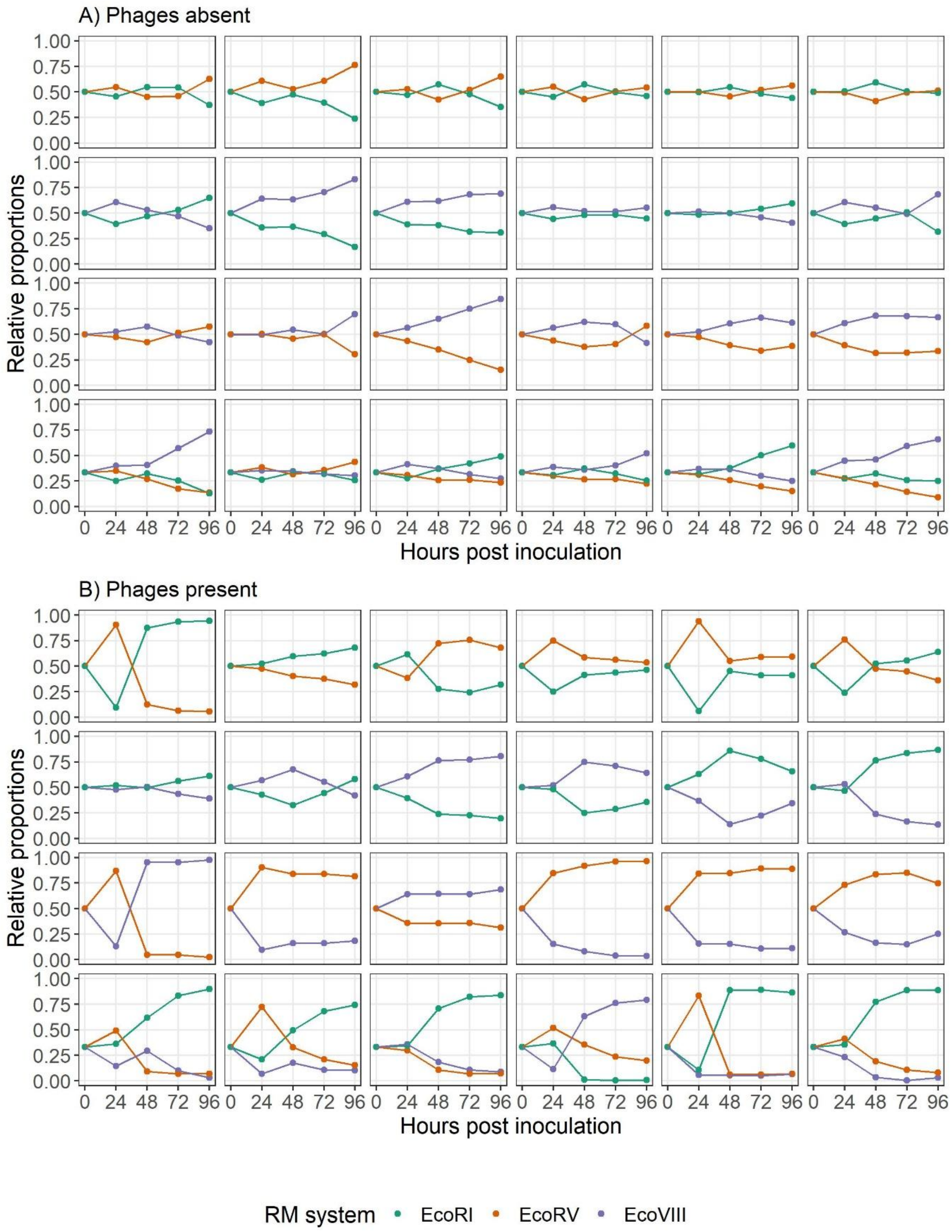


**Figure S4.** The relative frequencies (%) of strains carrying different RM systems (RM_type) over time, when phages are absent (A) or present (B). Different panels denote different biological replicates.

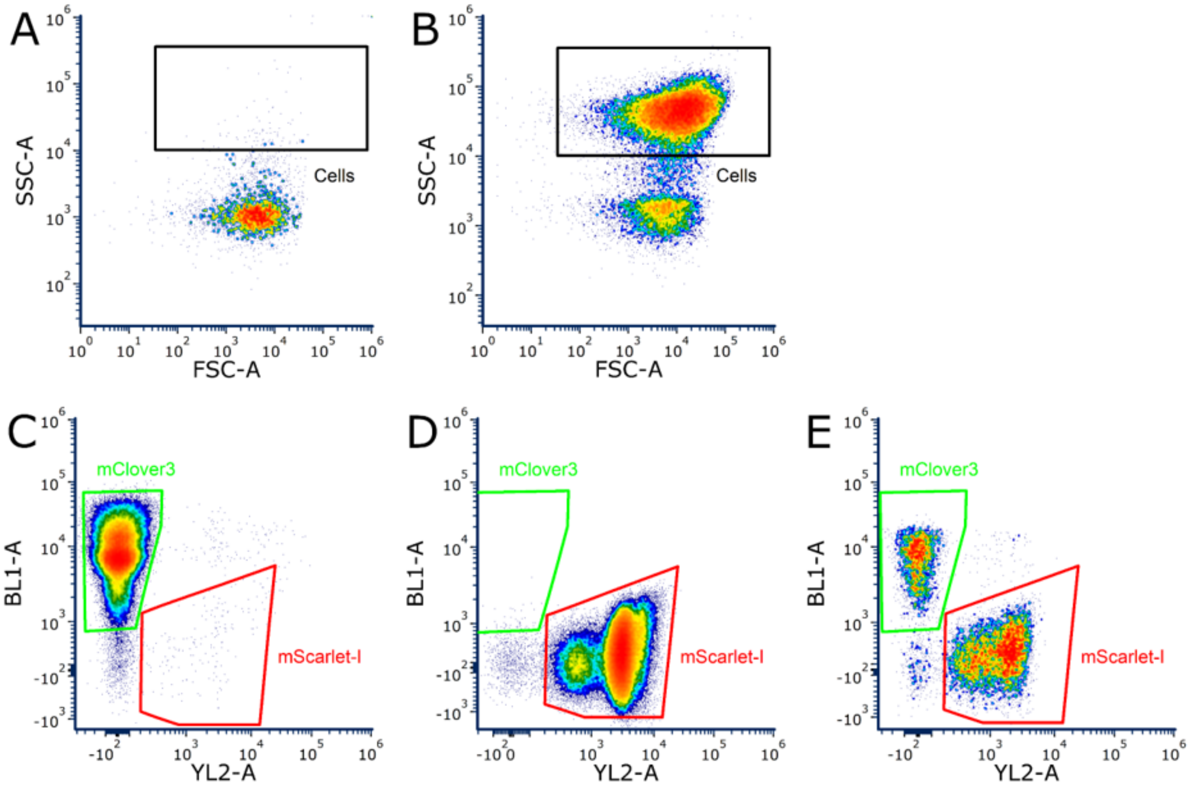


**Figure S5. Flow cytometry gating strategy for two-fluorophore discrimination using the Attune NxT.** Representative gating plots illustrating the analysis pipeline used to distinguish mClover3 and mScarlet-I expressing cells. (A) Forward scatter (FSC-A) vs. side scatter (SSC-A) plot of a cell-free control sample containing only phosphate-buffered saline (PBS) and λvir. (B) FSC-A vs. SSC-A plot of a cell-containing sample, showing separation of the cell population from debris. (C) BL1-A vs. YL2-A fluorescence plot showing the gating of mClover3-expressing cells (only plotting events gated as Cells from B). (D) BL1-A vs. YL2-A fluorescence plot showing the gating of mScarlet-I-expressing cells (only plotting events gated as Cells from B); note there is some bimodality across the YL2-A axis, however, both sub-populations of mScarlet-I-expressing cells remain distinguishable from mClover3-expressing cells. (E) BL1-A vs. YL2-A fluorescence plot of a mixed mClover3- and mScarlet-I-expressing population at 96 hours, confirming that both fluorescent reporters are distinguishable and stably maintained throughout the duration of the experiment.

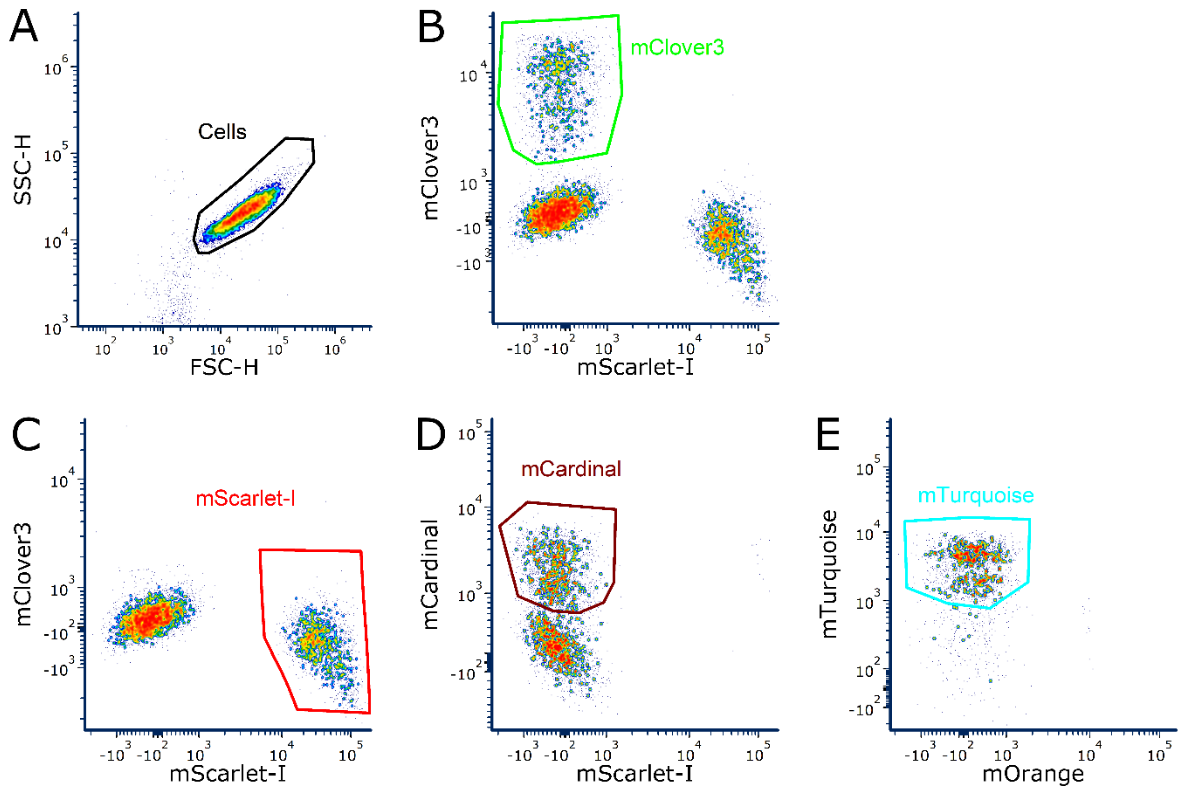


**Figure S6. Flow cytometry gating strategy for four-fluorophore discrimination using the Cytek Aurora.** Representative gating plots illustrating the sequential gating pipeline used to distinguish mClover3-, mScarlet-I-, mCardinal-, and mTurquoise2-expressing cells. All panels plot data from the same biological replicate. (A) FSC vs. SSC plot showing separation of the cell population from debris. (B) mScarlet-I vs. mClover3 fluorescence plot showing the gating of mClover3-expressing cells; ungated events are passed to (C). (C) mScarlet-I vs. mClover3 fluorescence plot showing the gating of mScarlet-I-expressing cells from the ungated events of (B); ungated events are passed to (D). (D) mScarlet-I vs. mCardinal fluorescence plot showing the gating of mCardinal-expressing cells from the ungated events of (C); ungated events are passed to (E). (E) mOrange vs. mTurquoise2 fluorescence plot showing the gating of mTurquoise2-expressing cells from the ungated events of (D).
